# AAV9-mediated βIII-tubulin Ser172 phospho-mimic expression improves arrhythmic and inflammatory remodeling in dystrophic cardiomyopathy

**DOI:** 10.64898/2026.08.04.742904

**Authors:** Delong Zhou, Vasisht Yegneshwaran, Nehal Kamal Ali, Geovanni Geukgeuzian, Elam Mesa, Lai-Hua Xie, Diego Fraidenraich

**Author notes:** Corresponding author. 185 South Orange Avenue, Medical Sciences Building G-667, Newark, NJ, 07103, USA.

## Abstract

**Background:** Duchenne muscular dystrophy (DMD) cardiomyopathy is characterized by progressive microtubule remodeling, connexin-43 (Cx43) dysregulation, and ventricular arrhythmias. We previously demonstrated phospho-mimic knock-in of βIII-tubulin S172E preserves microtubule organization and attenuates cardiac pathology in mdx mice. However, whether these protective effects can be reproduced using a clinically relevant gene-delivery strategy remains unknown.

**Methods and Results:** We generated a cardiomyocyte-specific adeno-associated virus serotype 9 (AAV9) vector expressing phospho-mimic βIII-tubulin (Tubb3-S172E) under the cardiac troponin T promoter and delivered it to 4-5-month-old wild-type and mdx mice. Cardiac Tubb3-S172E expression was confirmed by quantitative qPCR and immunoblotting. In mdx mice, AAV9-mediated Tubb3-S172E expression significantly reduced mononuclear inflammatory infiltration, restored Cx43 localization at intercalated discs, and attenuated isoproterenol-induced arrhythmia susceptibility. In contrast, cardiac fibrosis, Nav1.5 protein expression, and peak sodium current density were not significantly improved. Overexpression of wild-type βIII-tubulin in healthy hearts increased Cx43 lateralization and arrhythmia susceptibility, indicating that βIII-tubulin phosphorylation state rather than protein abundance determines its protective function.

**Conclusions:** Cardiomyocyte-targeted delivery of phospho-mimic βIII-tubulin partially recapitulates the protective effects observed in the genetic S172E knock-in model. These findings identify βIII-tubulin Ser172 phosphorylation as a critical regulator of microtubule-dependent electrical remodeling and support therapeutic modulation of this pathway in Duchenne muscular dystrophy cardiomyopathy.

**Research Perspective:**

- Cardiomyocyte-targeted AAV9 delivery of phospho-mimic âIII-tubulin improves Cx43 organization, inflammatory remodeling, and arrhythmia susceptibility in dystrophic hearts, demonstrating that therapeutic modulation of βIII-tubulin Ser172 phosphorylation partially recapitulates the protective effects observed in the genetic S172E model.
- The dissociation between improved electrical remodeling and persistent Nav1.5 and fibrotic abnormalities suggests that βIII-tubulin Ser172 phosphorylation selectively regulates specific microtubule-dependent pathological pathways in dystrophic cardiomyopathy.
- Future studies should define the molecular mechanisms linking βIII-tubulin Ser172 phosphorylation to cardiomyocyte-immune cell communication and determine how this pathway coordinates electrical and inflammatory remodeling in dystrophic hearts.

## Introduction

Duchenne muscular dystrophy (DMD)-associated cardiomyopathy is a leading cause of mortality in patients with DMD and is characterized by progressive myocardial fibrosis, ventricular contractile dysfunction, and an increased susceptibility to arrhythmias [1–3]. As a central component of the dystrophin-glycoprotein complex, dystrophin mechanically links the intracellular cytoskeleton to the extracellular matrix [4]. Loss of dystrophin therefore compromises sarcolemmal stability and promotes remodeling of cytoskeletal and intercellular junctional networks in DMD cardiomyocytes [5].

Microtubules are essential components of the cardiomyocyte cytoskeleton and regulate intracellular signaling, membrane-protein trafficking, organelle positioning, and the maintenance of cardiomyocyte architecture [6–8]. Increasing evidence implicates microtubule remodeling in the pathogenesis of DMD cardiomyopathy, linking dystrophin deficiency to altered tubulin post-translational modifications (PTM) [9], impaired cytoskeletal mechanotransduction [10, 11], disrupted connexin-43 and ion-channel homeostasis [12, 13], and subsequent electrical and structural cardiac remodeling [14].

Connexin-43 (Cx43) is the predominant ventricular gap-junction protein and supports electrical and metabolic coupling between cardiomyocytes [15, 16]. In DMD hearts, Cx43 undergoes pathological remodeling characterized by altered phosphorylation, increased protein abundance, and redistribution from the intercalated discs to the lateral sarcolemma (lateralization) [12, 13, 17]. This remodeling is associated with oxidative stress, aberrant Cx43 hemichannel activity, and increased susceptibility to arrhythmias [13, 17–19]. Accordingly, normalization of Cx43 abundance and localization, as well as inhibition of pathological hemichannel activity, has been proposed as a therapeutic strategy for DMD cardiomyopathy [13, 19, 20].

Our previous studies identified a reciprocal relationship between Cx43 remodeling and microtubule organization in DMD cardiomyopathy. Phospho-mimic substitution of the Cx43 C-terminal serine triplet (Ser325/328/330, Cx43-S3E) normalized Cx43 remodeling in mdx hearts and reduced pathological microtubule accumulation and orthogonal reorganization [12, 13]. Conversely, colchicine-mediated microtubule modulation reduced microtubule remodeling and cardiac fibrosis while restoring Cx43 localization at the intercalated discs in mdx hearts [12]. These findings support a functional regulatory link between the microtubule cytoskeleton and sarcolemmal junctional proteins. Consistent with this model, our recent study demonstrated that phospho-mimic substitution of βIII-tubulin Ser172 (S172E) restored microtubule organization and dynamics in mdx hearts, accompanied by improved Cx43 phosphorylation and reduced lateralization [14].

To determine whether these protective effects could be reproduced without germline genetic modification, we developed an AAV9-based approach to express phospho-mimic βIII-tubulin (Tubb3-S172E), under the control of the cardiac troponin T (cTnT) promoter. This strategy enabled us to test whether cardiomyocyte-targeted expression of exogenous Tubb3-S172E is sufficient to ameliorate cardiac abnormalities in mdx mice. We found that cardiomyocyte-specific expression of phospho-mimic βIII-tubulin reduced mononuclear infiltration, improved Cx43 localization, and attenuated isoproterenol-induced arrhythmia susceptibility, despite having limited effects on Nav1.5 remodeling and cardiac fibrosis. This study not only provides proof-of-concept that therapeutic delivery of phospho-mimic βIII-tubulin can partially recapitulate the protective effects observed in the S172E knock-in model, but also further establishes βIII-tubulin Ser172 phosphorylation as a key regulator of microtubule-dependent electrical remodeling and a potential therapeutic target for Duchenne muscular dystrophy cardiomyopathy.

## Results

### AAV9-mediated cardiomyocyte-targeted expression of phospho-mimic βIII-tubulin in WT and mdx mice

The AAV9 vector was designed to deliver mouse Tubb3, encoding either wild-type βIII-tubulin or the phospho-mimic S172E mutant, under the control of the cardiac troponin T (cTnT) promoter to enhance cardiomyocyte-specific expression. A P2A-linked EGFP reporter was included downstream of Tubb3 to facilitate assessment of viral transduction (Figure 1A). WT and mdx mice received a single retro-orbital injection of PBS, AAV9-mTubb3-WT, or AAV9-mTubb3-S172E, and ventricular tissues were collected 21 days after administration.

**Figure 1.**
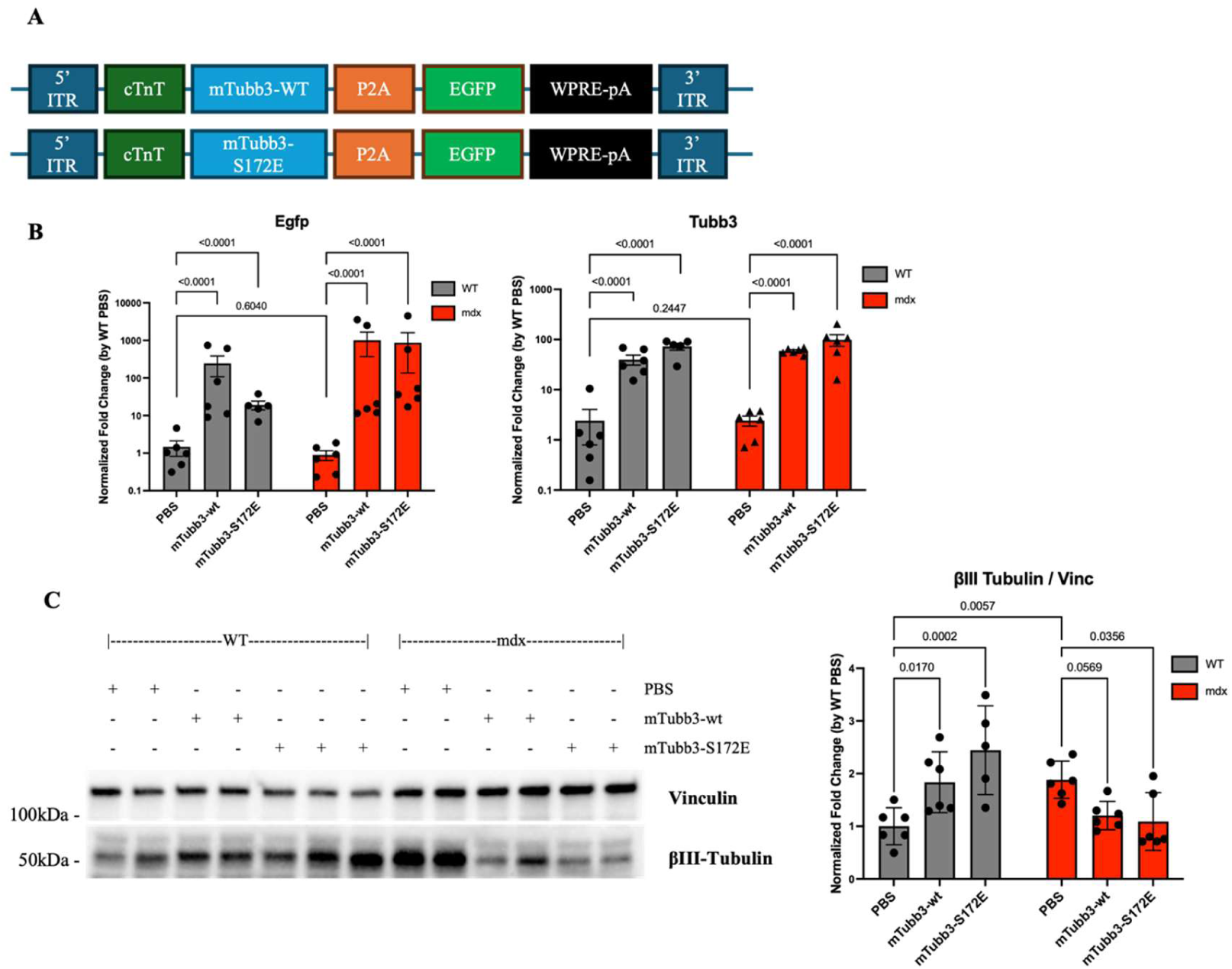
AAV9 vector design and validation of cardiac βIII-tubulin overexpression. **(A)** Schematic representation of the AAV9 construct used for cardiac gene delivery. The vector contains a cardiac troponin T promoter (cTnT) driving expression of either mouse wild-type Tubb3 or phospho-mimic Tubb3-S172E, followed by a P2A-linked EGFP reporter. **(B)** Quantitative PCR analysis of total *Tubb3* and *Egfp* mRNA expression across experimental groups. Expression was normalized to the average of the housekeeping genes *Rps18* and *Hmbs* and expressed as fold change relative to the WT + PBS group. The *Tubb3* primer set detects endogenous and exogenous *Tubb3* transcripts, including the *Tubb3-S172E* phospho-mimic transcript. Data are shown on a log10 scale. **(C)** Representative Western blot and quantification of βIII-tubulin protein expression in ventricular tissues across experimental groups, normalized to vinculin. Data are presented as mean ± SEM. Sample size was n=6 for each group, except for WT mice treated with AAV9-Tubb3-S172E, where n=5. Statistical analysis was performed using two-way ANOVA followed by Holm-Sidak post hoc multiple-comparisons test. Exact P values are indicated in the figure.

Because native EGFP fluorescence was relatively weak in cardiac sections and partially limited by tissue autofluorescence, viral transduction was confirmed using complementary approaches. Quantitative PCR analysis of ventricular RNA showed increased *Egfp* and *Tubb3* mRNA levels following AAV9 treatment (Figure 1B). In addition, anti-EGFP immunofluorescence detected positive signal in cardiac sections using the Alexa Fluor 555 channel, further supporting successful cardiac transduction (Supp. Figure 1A). At the protein level, AAV9-mTubb3-WT and AAV9-mTubb3-S172E increased total βIII-tubulin abundance in WT hearts. In contrast, despite increased *Tubb3* mRNA expression following AAV9 administration, βIII-tubulin protein levels in mdx hearts showed a decreasing trend after treatment.

Gross physiological parameters were assessed by measuring body and tissue weights normalized to tibia length (Supp. Figure 1B). At 4-5 months of age, mdx mice exhibited higher normalized liver, heart, and body weights than WT mice, consistent with broad disease-associated changes in tissue growth and remodeling. Normalized liver weight was also evaluated because the liver is a major site of systemic AAV9 biodistribution and therefore represents a relevant organ for evaluating potential off-target effects. Neither AAV9-mTubb3-WT nor AAV9-mTubb3-S172E produced detectable changes in normalized body, heart, or liver weight in WT or mdx mice, indicating that AAV9-mediated *Tubb3* expression was not associated with overt alterations in these gross physiological parameters.

### Phospho-mimic βIII-tubulin attenuates cardiac inflammatory remodeling in mdx mice

Following confirmation of successful cardiac transduction and transgene expression, hematoxylin and eosin (H&E) staining was performed to assess the degree of mononuclear infiltration. Compared with PBS-treated WT mice, PBS-treated mdx hearts at 4-5 months of age exhibited a marked accumulation of mononuclear cells within the ventricular myocardium, indicative of pronounced inflammatory cell infiltration (Figure 2A). Administration of either AAV9-mTubb3-WT or AAV9-mTubb3-S172E reduced mononuclear infiltration in mdx hearts, with the phospho-mimic S172E treatment producing the greatest effect and bringing mononuclear infiltration to levels comparable to those observed in WT hearts (Figure 2B). In contrast, neither AAV9-mTubb3-WT nor AAV9-mTubb3-S172E altered mononuclear infiltration in WT mice.

**Figure 2.**
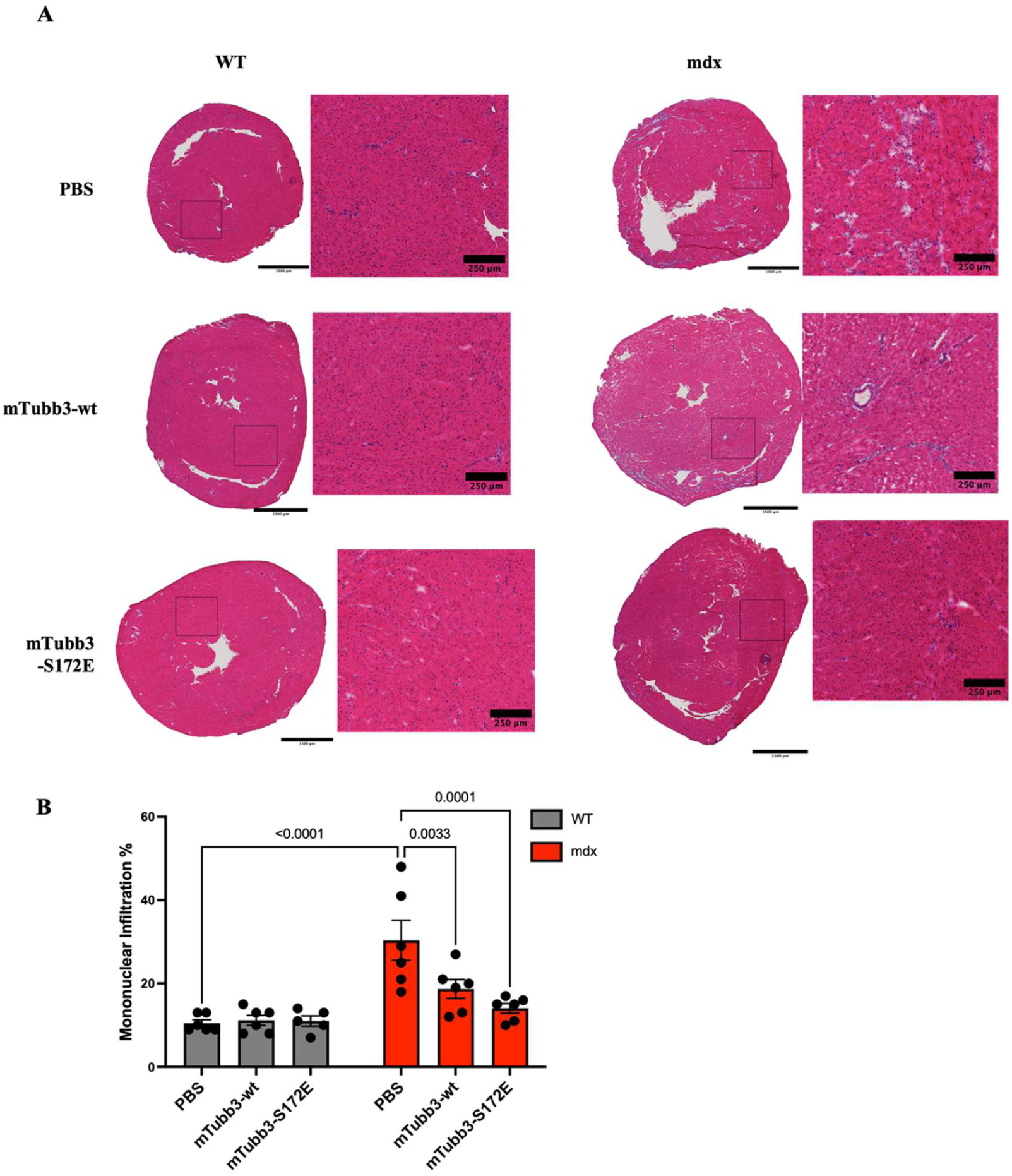
Cardiac expression of WT or phospho-mimic S172E βIII-tubulin reduces mononuclear infiltration in mdx hearts. **(A)** Representative hematoxylin and eosin (H&E)-stained ventricular sections from WT and mdx mice treated with PBS, AAV9-mTubb3-WT, or AAV9-mTubb3-S172E at 4-5 months of age. Higher-magnification insets highlight regions of mononuclear infiltration. **(B)** Mononuclear infiltration was quantified using a preset grid-overlay method and expressed as the percentage of grid regions positive for mononuclear infiltration relative to the total number of grid regions analyzed. Scale bars: 1500 μm for low-magnification images and 250 μm for higher-magnification insets. Data are presented as mean ± SEM. Sample size was n=6 for each group, except for WT mice treated with AAV9-Tubb3-S172E, where n=5. Statistical analysis was performed using two-way ANOVA followed by Holm-Sidak post hoc multiple-comparisons test. Exact P values are indicated in the figure.

Picrosirius Red (PSR) staining demonstrated increased interstitial and perivascular fibrosis in PBS-treated mdx hearts compared with PBS-treated WT hearts (Supp. Figure 2A). However, unlike the reduction in mononuclear infiltration, neither AAV9-mTubb3-WT nor AAV9-mTubb3-S172E significantly altered the extent of cardiac fibrosis in mdx mice (Supp. Figure 2B). The response was largely restricted to inflammatory remodeling, with little effect on established fibrosis in mdx hearts.

### Phospho-mimic βIII-tubulin restores Cx43 localization at the intercalated discs in mdx hearts

To assess whether AAV9-mediated expression of phospho-mimic βIII-tubulin influences Cx43 localization in the dystrophic heart, immunofluorescence staining was performed to assess the distribution of Cx43 relative to the intercalated disc marker N-cadherin. In ventricular cryosections, N-cadherin was labeled with Alexa Fluor 555 (red) and Cx43 with Alexa Fluor 488 (green), with overlapping signals appearing yellow (Figure 3A). Compared with PBS-treated WT hearts, in which Cx43 was predominantly localized to N-cadherin-positive intercalated discs, PBS-treated mdx hearts exhibited reduced colocalization of Cx43 with N-cadherin and increased Cx43 signal along the lateral membrane. AAV9-mediated expression of phospho-mimic βIII-tubulin (AAV9-mTubb3-S172E) markedly increased Cx43 colocalization with N-cadherin, restoring the overlap ratio to a level comparable to that observed in WT hearts (Figure 3B).

**Figure 3.**
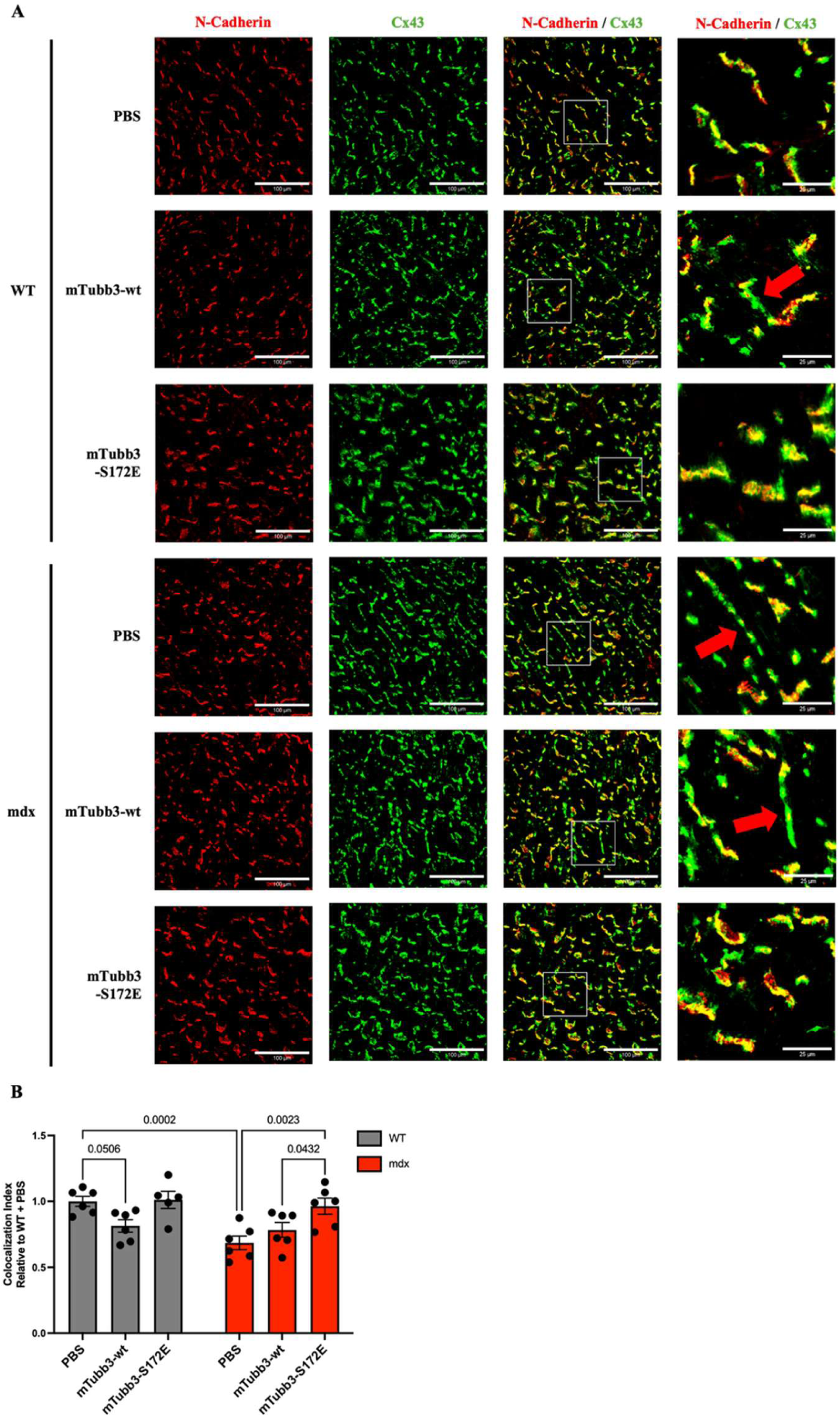
Cardiac expression of phospho-mimic S172E βIII-tubulin normalizes the Cx43 lateralization in mdx hearts. **(A)** Representative confocal images of ventricular cryosections co-stained for Cx43 (green) and N-Cadherin (red). Insets highlight representative lateralized Cx43 (red arrows). Scale bar: 100 µm (main) and 25 µm (insets). **(B)** Quantification of the Cx43/N-Cadherin colocalization index (fraction of colocalized Cx43 to total Cx43, normalized to the WT + PBS group). Data are presented as mean ± SEM. Sample size was n=6 for each group, except for WT mice treated with AAV9-Tubb3-S172E, where n=5. Statistical analysis was performed using two-way ANOVA followed by Holm-Sidak post hoc multiple-comparisons test. Exact P values are indicated in the figure.

Notably, AAV9-mediated overexpression of WT βIII-tubulin in WT hearts reduced Cx43 colocalization with N-cadherin and increased Cx43 lateralization (Figure 3B). These observations demonstrate that phospho-mimic βIII-tubulin improves Cx43 localization in mdx hearts while WT βIII-tubulin overexpression alters Cx43 distribution in WT hearts.

### Phospho-mimic βIII-tubulin is associated with reduced arrhythmia susceptibility in mdx mice following isoproterenol challenge

As Cx43 lateralization has been closely linked to ventricular arrhythmogenesis, electrocardiography was performed to evaluate arrhythmia susceptibility following AAV administration. Mice were anesthetized with Avertin and challenged with isoproterenol (ISO), after which ECG recordings were obtained for 1 hour. Compared with PBS-treated WT mice, which exhibited the expected ISO-induced sinus tachycardia with occasional single premature ventricular contractions (PVCs), PBS-treated mdx mice developed frequent PVCs and episodes of nonsustained ventricular tachycardia (VT) (Figure 4A). AAV9-mediated expression of phospho-mimic βIII-tubulin was associated with a lower arrhythmia severity score in mdx mice compared with PBS-treated mdx controls (Figure 4B). Although this reduction did not reach statistical significance after multiple-comparison correction (P < 0.08), the distribution of individual arrhythmia scores largely overlapped with those observed in PBS-treated WT mice (Figure 4B).

**Figure 4.**
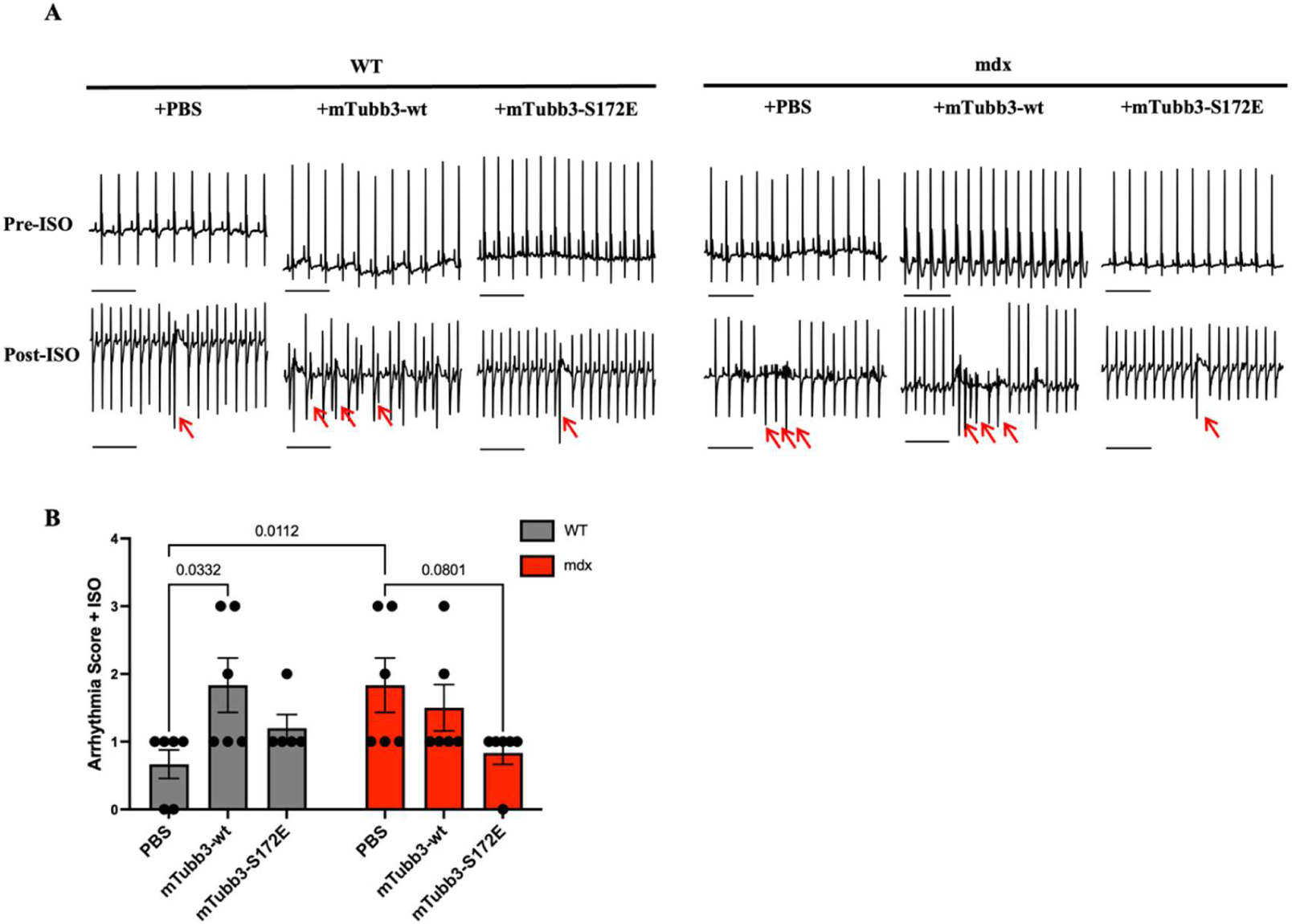
AAV9-mediated expression of phospho-mimic S172E βIII-tubulin is associated with reduced isoproterenol-induced arrhythmia severity in mdx mice. **(A)** Representative ECG tracings from WT and mdx mice treated with PBS, AAV9-mTubb3-WT, or AAV9-mTubb3-S172E at 4-5 months of age before and after isoproterenol administration. Arrows indicate premature ventricular complexes (PVCs). Representative tracings from WT mice treated with AAV9-mTubb3-WT and mdx mice treated with PBS or AAV9-mTubb3-WT show episodes of nonsustained ventricular tachycardia (VT) following isoproterenol administration. Scale bar: 500 ms. **(B)** Quantification of arrhythmia severity scores after isoproterenol administration. Arrhythmia severity was scored as follows: 0, no events; 1, single PVC; 2, paired PVCs; 3, multiple PVCs or nonsustained VT; 4, sustained VT; and 5, death due to asystole or ventricular fibrillation. Data are presented as mean ± SEM. Sample size was n=6 for each group, except for WT mice treated with AAV9-Tubb3-S172E, where n=5. Statistical analysis was performed using two-way ANOVA followed by Holm-Sidak post hoc multiple-comparisons test. Exact P values are indicated in the figure.

Notably, AAV9-mediated overexpression of WT βIII-tubulin in WT hearts increased arrhythmia severity following ISO challenge (Figure 4B). This finding is consistent with the increased Cx43 lateralization observed in WT hearts after AAV9-mTubb3-WT administration (Figure 3). Overall, the electrophysiological findings are consistent with improved ventricular electrical stability following phospho-mimic βIII-tubulin expression in mdx mice, while revealing distinct consequences of WT βIII-tubulin overexpression in healthy myocardium.

### Phospho-mimic βIII-tubulin does not restore Nav1.5 expression or sodium current density in mdx cardiomyocytes

Given the close functional interplay between Cx43 and Nav1.5 at the intercalated disc, we next examined whether phospho-mimic βIII-tubulin also influences Nav1.5 expression and sodium channel function. Western blot analysis demonstrated that Nav1.5 protein expression was significantly reduced in PBS-treated mdx hearts compared with PBS-treated WT hearts (Figure 5A). However, neither AAV9-mediated WT βIII-tubulin nor phospho-mimic βIII-tubulin expression altered Nav1.5 protein abundance in either WT or mdx hearts.

**Figure 5.**
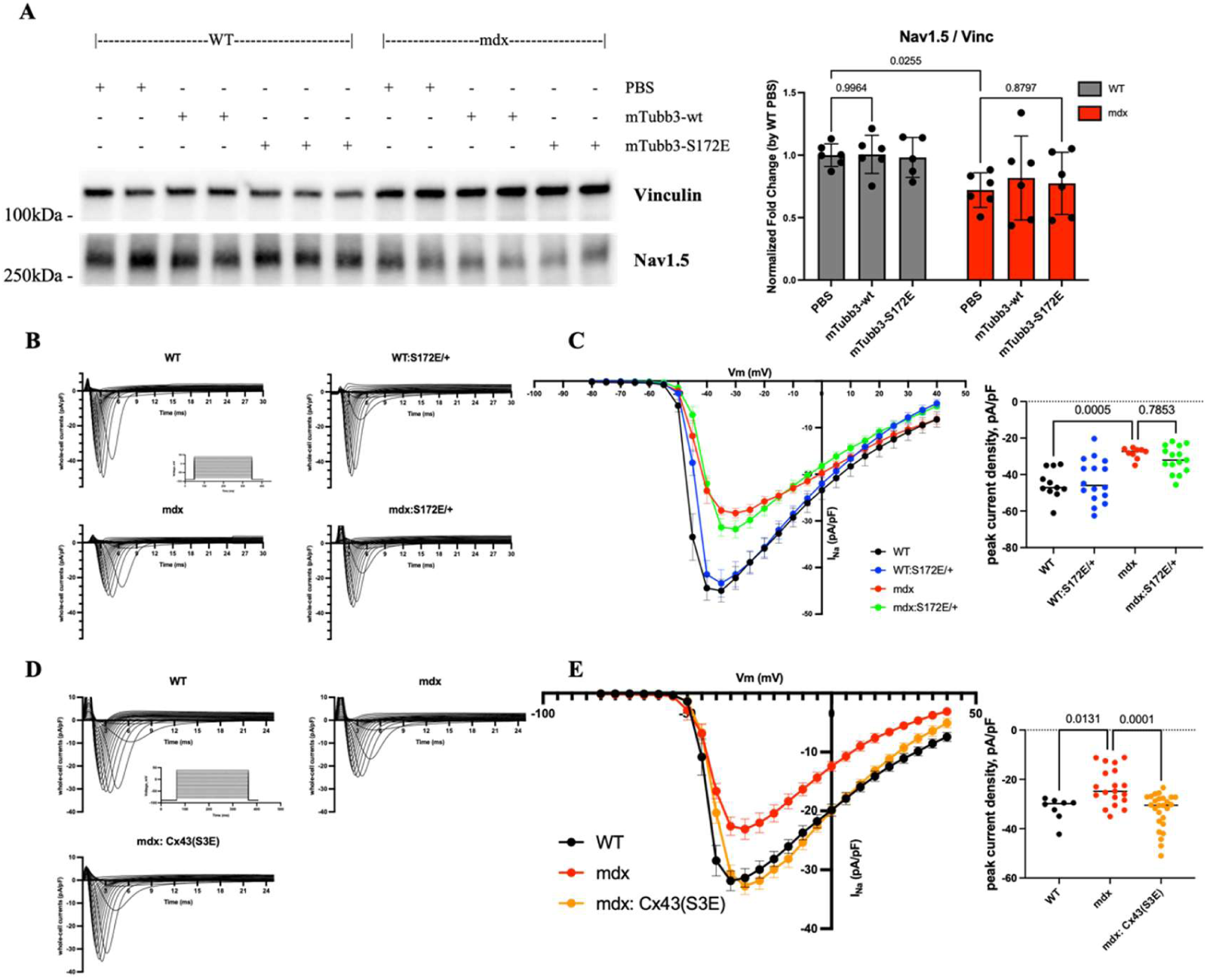
βIII-tubulin S172E does not restore Nav1.5 expression or sodium current density in mdx cardiomyocytes. **(A)** Representative Western blot images and quantification of Nav1.5 protein expression across experimental groups. The vinculin loading control is the same as that shown in Figure 1C, as they were derived from the same membrane. **(B)** Representative whole-cell sodium current (INa) traces recorded from cardiomyocytes isolated from 4–6-month-old WT, mdx, WT:S172E/+, and mdx:S172E/+ mice. **(C)** Normalized current density–voltage (I–V) relationships and quantification of peak INa density, showing reduced INa density in dystrophin-deficient cardiomyocytes that was not corrected by the S172E/+ mutation. Sample sizes were n=11 cells from N=4 mice for WT, n=16 cells from N=5 mice for WT:S172E/+, n=9 cells from N=4 mice for mdx, and n=15 cells from N=5 mice for mdx:S172E/+. **(D)** Representative whole-cell INa traces recorded from WT, mdx, and mdx:Cx43-S3E cardiomyocytes isolated from 3–5-month-old mice. **(E)** Normalized I–V relationships and quantification of peak INa density in WT, mdx, and mdx:Cx43-S3E cardiomyocytes, showing recovery of sodium current density in mdx:Cx43-S3E cardiomyocytes relative to mdx controls. Sample sizes were n=8 cells from N=2 mice for WT, n=19 cells from N=4 mice for mdx, and n=26 cells from N=4 mice for mdx:Cx43-S3E. Data are presented as mean ± SEM. For panel A, sample size was n=6 for each group, except for WT mice treated with AAV9-Tubb3-S172E, where n=5. Statistical analysis was performed using two-way ANOVA followed by Holm-Sidak post hoc multiple-comparisons test for panels A, with exact P values indicated in the figure. One-way ANOVA followed by Tukey post hoc multiple-comparisons test was used for panels C and E. *P<0.05, ***P<0.001.

To investigate whether phospho-mimic βIII-tubulin also restores sodium channel function, whole-cell patch-clamp recordings were performed in ventricular cardiomyocytes isolated from Tubb3-S172E/+ mice, a model previously shown to improve microtubule organization, Cx43 localization, and ventricular arrhythmia in mdx hearts [14]. Compared with WT cardiomyocytes, mdx cardiomyocytes exhibited a significant reduction in peak sodium current density (Figure 5B, C). However, the S172E/+ mutation did not restore peak sodium current density.

To further evaluate the relationship between Cx43 remodeling and sodium channel function, we next analyzed ventricular cardiomyocytes isolated from phospho-mimic Cx43-S3E mice, a genetic model previously shown to normalize Cx43 localization and reduce ventricular arrhythmias in mdx hearts [13]. Unlike the phospho-mimic βIII-tubulin S172E/+ mutation, phospho-mimic Cx43 significantly restored peak sodium current density in mdx cardiomyocytes to levels comparable to WT controls (Figure 5D, E).

Taken together, these findings indicate that although phospho-mimic βIII-tubulin improves Cx43 localization and ventricular electrical stability in mdx mice, these protective effects are not accompanied by restoration of Nav1.5 expression or sodium current density. In contrast, direct phospho-mimic modification of Cx43 was sufficient to normalize sodium current density, suggesting that recovery of sodium channel function is more closely associated with Cx43 remodeling than with phospho-mimic βIII-tubulin expression.

## Discussion

Microtubule remodeling is a well-recognized hallmark of cardiomyopathy progression and is characterized by altered post-translational modifications (PTMs) together with increased microtubule density and network disorganization [21–24]. Beyond providing structural support, the microtubule cytoskeleton regulates intracellular trafficking, organelle positioning, and multiple signaling pathways that are essential for cardiomyocyte homeostasis. In dystrophic hearts, increased detyrosination of α-tubulin promotes microtubule densification, enhances X-ROS signaling, and contributes to calcium dysregulation, ventricular arrhythmias, and contractile dysfunction [9, 10, 25]. Microtubule remodeling also disrupts T-tubule organization and sodium channel function, thereby contributing to impaired excitation-contraction coupling, calcium dysregulation, ventricular arrhythmogenesis, and contractile dysfunction [9, 11]. Current evidence indicates that individual microtubule PTMs regulate distinct aspects of cardiomyocyte biology. This raises the possibility that targeting specific PTMs, rather than globally altering microtubule abundance, may provide a more selective therapeutic strategy [26].

Beyond the well-characterized PTMs of α-tubulin, our previous study using phospho-mimic βIII-tubulin S172E knock-in mice demonstrated that reduced Ser172 phosphorylation contributes to microtubule disorganization, Cx43 remodeling, and ventricular arrhythmogenesis in mdx hearts [14]. These findings suggested that dysregulation of β-tubulin PTMs may represent an additional mechanism underlying microtubule remodeling in dystrophic cardiomyopathy. Because βIII-tubulin is a relatively low-abundance β-tubulin isoform in the heart, and the global S172E knock-in model may also affect non-cardiomyocyte cell types, we employed AAV9-mediated cardiomyocyte-specific expression of phospho-mimic βIII-tubulin driven by the cTnT promoter to determine its cell-specific role in the dystrophic myocardium (Fig. 1A). 21 days after a single retro-orbital injection, successful cardiac transduction was confirmed by increased ventricular *Tubb3* and *Egfp* mRNA expression together with positive anti-EGFP immunofluorescence staining (Fig. 1B; Supp. Fig. 1A).

AAV-mediated *Tubb3* overexpression increased βIII-tubulin protein abundance in WT hearts as expected. By contrast, successful transgene expression at the mRNA level was not accompanied by a corresponding increase in βIII-tubulin protein in mdx hearts (Figure 1C). Given that dystrophic hearts already exhibit elevated endogenous βIII-tubulin and pan-β-tubulin expression in both mice and human DMD myocardium [14], this differential response suggests that βIII-tubulin protein abundance is regulated differently in the dystrophic myocardium through mechanisms beyond transcriptional control. One explanation is that dystrophic myocardium limits further βIII-tubulin accumulation through tubulin autoregulation, a feedback mechanism that restricts excess tubulin accumulation and preserves cytoskeletal homeostasis [27–29]. In addition, extensive microtubule remodeling in mdx hearts may impair incorporation or stability of exogenous βIII-tubulin within the microtubule lattice. The dissociation between transcript and protein abundance suggests the existence of disease-specific post-translational regulatory mechanisms in dystrophic myocardium.

Notably, neither WT nor phospho-mimic S172E βIII-tubulin expression altered the already low level of mononuclear infiltration in WT hearts. In contrast, both AAV9-mTubb3-WT and AAV9-mTubb3-S172E significantly reduced mononuclear infiltration in mdx hearts, with the greatest improvement observed following S172E expression (Figure 2), despite the absence of an overall increase in βIII-tubulin protein abundance. The genotype dependence probably reflects the low baseline inflammatory burden in WT hearts, leaving little opportunity for further improvement. In contrast, modulation of the remodeled microtubule network in mdx hearts may reduce cardiomyocyte injury and secondary inflammatory-cell recruitment [25, 30]. Although immunoblotting could not distinguish endogenous from AAV-derived βIII-tubulin, the greater benefit observed with S172E supports the concept that phosphorylation state of βIII-tubulin, rather than its protein abundance, determines its protective effect [26, 31].

Pathological lateralization of connexin 43 (Cx43) is a well-established hallmark of DMD cardiomyopathy and contributes to ventricular arrhythmogenesis [13, 18]. Proper trafficking of Cx43 to the intercalated disc depends on an intact and properly organized microtubule network [32]. In our previous study, we demonstrated that phospho-mimic βIII-tubulin S172E restored microtubule organization, normalized Cx43 localization, and reduced isoproterenol (ISO)-induced arrhythmias in mdx mice [14]. The present study extends these findings by demonstrating that cardiomyocyte-specific AAV9-mediated expression of phospho-mimic βIII-tubulin S172E similarly reduced Cx43 lateralization and improved its localization at the intercalated discs. These structural improvements were accompanied by reduced susceptibility to isoproterenol-induced ventricular arrhythmias, whereas WT βIII-tubulin overexpression failed to produce comparable benefits (Figures 3 and 4). These findings further support the concept that the biological activity of βIII-tubulin is determined predominantly by its phosphorylation state rather than its abundance. Although both constructs expressed the same βIII-tubulin isoform, only phospho-mimic S172E restored Cx43 localization and reduced arrhythmia susceptibility, indicating that Ser172 phosphorylation is required for the protective effects of βIII-tubulin on microtubule-dependent Cx43 trafficking. Conversely, WT βIII-tubulin overexpression further increased Cx43 lateralization and arrhythmia susceptibility in healthy hearts. This observation is consistent with our previous finding that overexpression of βIVb-tubulin disrupted microtubule directionality in WT myocardium [14], suggesting that excessive β-tubulin expression perturbs the highly organized microtubule architecture of healthy cardiomyocytes, thereby impairing membrane protein trafficking and reducing electrical stability.

Although phospho-mimic S172E reduced mononuclear infiltration, it did not attenuate established cardiac fibrosis in mdx hearts (Supp. Figure 2). This difference likely reflects the distinct time courses of inflammatory and fibrotic remodeling [30]. AAV treatment was initiated at 4-5-months of age and analyzed only 21 days later, a period that may be sufficient to suppress ongoing inflammation but insufficient to reverse established extracellular matrix remodeling [33]. Consistent with this interpretation, fibrosis was reduced only in aged mdx:S172E/+ knock-in mice in our previous study [14]. βIII-tubulin phosphorylation therefore appears to preferentially influence early cardiomyocyte-dependent remodeling, whereas established fibrosis is likely to require longer treatment or additional therapeutic strategies.

Given the well-established structural and functional relationship between Cx43 and Nav1.5 at the intercalated disc [34, 35], we next investigated whether restoration of Cx43 remodeling by phospho-mimic βIII-tubulin was accompanied by recovery of Nav1.5. Despite improving Cx43 localization and reducing arrhythmia susceptibility, phospho-mimic βIII-tubulin failed to restore either Nav1.5 protein expression or peak sodium current density in mdx cardiomyocytes (Figure 5A-C). In contrast, our previous Cx43-S3E phospho-mimic model restored sodium current density (Figure 5D-E), and was previously shown to produce a comparable improvement in Cx43 remodeling [13]. These findings suggest that restoration of microtubule organization and Cx43 localization alone is insufficient to fully normalize Nav1.5 remodeling. Instead, complete restoration of Nav1.5 remodeling likely requires additional molecular mechanisms beyond normalization of the microtubule network. Importantly, the improvement in arrhythmia susceptibility despite persistent Nav1.5 remodeling indicates that complete recovery of Nav1.5 is not required for the anti-arrhythmic effects of βIII-tubulin Ser172 phosphorylation.

This study has limitations. First, although phospho-mimic βIII-tubulin consistently reduced arrhythmia severity in mdx mice, the difference did not reach statistical significance, warranting validation in larger cohorts. Second, while cardiomyocyte-specific βIII-tubulin modification reduced mononuclear infiltration, the mechanisms linking cardiomyocyte remodeling to immune-cell recruitment remain unclear. Given the cardiomyocyte-specific expression driven by the cTnT promoter, these anti-inflammatory effects are likely mediated through indirect cardiomyocyte-non-cardiomyocyte communication, which requires further investigation.

In summary, the present study demonstrates that cardiomyocyte-specific expression of phospho-mimic βIII-tubulin S172E partially attenuates dystrophic cardiomyopathy, including mononuclear infiltration, Cx43 organization, and arrhythmia susceptibility. These findings further support the concept that the phosphorylation state of βIII-tubulin, rather than its abundance, is a critical determinant of its biological function in the dystrophic heart. Together with our previous genetic studies, this work identifies βIII-tubulin Ser172 phosphorylation as a promising therapeutic target for modulating microtubule remodeling and improving electrical stability in Duchenne muscular dystrophy cardiomyopathy.

## Materials and Methods

### Animals and AAV9 administration

Wild-type (WT, C57BL/6J) and dystrophin-deficient mdx (C57BL/10ScSn-Dmd/J) mice were obtained from The Jackson Laboratories (Bar Harbor, ME). The generation, breeding, and maintenance of phospho-mimic βIII-tubulin Ser172 mutant mice (Tubb3-S172E) [14], and phospho-mimic connexin-43 mutant mice carrying substitutions at Ser325/328/330 (Cx43-S3E) [12, 13], have been described previously. The WT and mdx mice used for AAV9 studies were at 4-5 months of age, Tubb3-S172E mutant mice at 4-6 months of age, and Cx43-S3E mice at 3-5 months of age. Both male and female mice were included because no sex-dependent differences were identified at this age in our previous studies [12–14]. Investigators were blinded to genotype and treatment during data acquisition and analysis. Sample sizes were selected based on the previous studies [12, 14] and prior experience with these experimental models. Sample sizes for individual experiments are reported in the corresponding figure legends.

Recombinant adeno-associated virus serotype 9 (AAV9) vectors expressing WT mouse *Tubb3* (AAV9-mTubb3-wt) or phospho-mimic mouse *Tubb3*-S172E (AAV9-mTubb3-S172E) were packaged by VectorBuilder. Expression was driven by the cardiac troponin T promoter. Viral preparations were diluted in sterile phosphate-buffered saline (PBS) and administered to mice at 3-4 months of age by retro-orbital injection at a dose of 5x10^11^ GC/mouse in a total volume of 100 µL. Mice were randomly assigned into different groups and anesthetized with 3% isoflurane during injection. Control mice received 100 µL PBS through the same route. Electrocardiography was performed 3 weeks after the single administration, after which the mice were euthanized for tissue collection.

For electrocardiographic recordings, prolonged anesthesia was induced by a single intraperitoneal injection of Avertin at 290 mg/kg (12.5 mg/mL of 2,2,2-tribromethanol with 2.5% (v/v) of 2-methyl-2-butanol). For tissue collection, mice were deeply anesthetized with isoflurane in a closed induction chamber and euthanized by cervical dislocation after the absence of a response to toe pinch was confirmed. All animal procedures were performed in accordance with the National Institutes of Health guidelines and were approved by the Institutional Animal Care and Use Committee (IACUC) of Rutgers New Jersey Medical School. All tissues were collected postmortem.

### Electrocardiography

The electrocardiography (ECG) was performed as previously described [12, 14]. Anesthetized mice were positioned on a circulating-water heating pad maintained at 37 °C. After a 5-minute baseline recording, isoproterenol (ISO, Sigma, 5 mg/kg, i.p.) was administered and electrocardiographic signals were recorded continuously for an additional 60 minutes. Arrhythmias were scored according to the maximum severity observed during the recording period [13, 14]: 0 = no arrhythmia; 1 = single premature ventricular contraction (PVC); 2 = paired PVCs; 3 = multiple PVCs or nonsustained ventricular tachycardia (VT); 4 = sustained VT, defined as more than 10 consecutive ventricular beats; 5 = death, defined by asystole or ventricular fibrillation. Recordings were analyzed using Clampfit software (Ver. 11.2.2.17). Arrhythmia scores were independently assigned by two investigators blinded to genotype and treatment.

### Quantitative real-time PCR analysis

Total RNA was isolated from ventricular tissues using TRIzol reagent (Invitrogen, 15596026) according to the manufacturer’s instructions. 1 µg of total RNA was reverse-transcribed in a final volume of 40 µL using the High-Capacity cDNA Reverse Transcription Kit (Applied Biosystems, 4368814) with a 1:1 equal mixture of random hexamers and oligo(dT)_16_ primers (Invitrogen N8080128). qPCR was performed in 10 µL reactions using PowerUp SYBR Green Master Mix (Applied Biosystems, A25741) on Bio-Rad CFX96 Real-Time PCR Detection System. The thermal cycling conditions were described previously [14]. Melt-curve analysis was performed after amplification to assess reaction specificity. The following primer sequences were used:

*Tubb3*: (F) CAGATAGGGGCCAAGTTCTGG; (R) GCAGGTCTGAGTCCCCTACA.

*Egfp*: (F) GACAAGCAGAAGAACGGCATC; (R) GCTCAGGTAGTGGTTGTCGG.

*Rps18*: (F) CCACTTTTGGGGCCTTCGT; (R) GCAAAGGCCCAGAGACTCATT.

*Hmbs*: (F) GTGCCTACCATACTACCTCCTG; (R) ACTCTCCTCAGAGAGCTGGTTC.

Relative transcript abundance was calculated using the 2^(-ΔΔCq) method [36], and the statistical analyses were performed using ΔCq values. Target-gene expression was normalized to the geometric mean of the reference genes *Rps18* and *Hmbs*, and is presented as fold change relative to the WT + PBS group.

### Western blot analysis

Snap-frozen ventricular tissue was homogenized in RIPA lysis buffer as previously described [14]. Protein samples were mixed with Laemmli buffer (Bio-Rad, 1610747) containing β-mercaptoethanol and denatured at 99 °C for 5 minutes. Equal amounts of protein were separated using 10% SDS-PAGE gels (Bio-Rad, 4561033) and transferred to nitrocellulose membranes. Membranes were blocked for 1 hour at room temperature in 5% blocking buffer (Bio-Rad, 1706404) prepared in PBS containing 0.1% (v/v) Tween-20. Membranes were then incubated with primary antibodies overnight at 4 °C and with HRP-conjugated secondary antibodies for 1 hour at room temperature. Chemiluminescent signals were developed using Clarity Max Western ECL substrate kit (Bio-Rad, 1705062), acquired and quantified using ImageLab (Bio-Rad). The following primary antibodies were used: Vinculin (Sigma V9131, 1:2000); βIII-tubulin (Abcam ab78078, 1:1000); Nav1.5 (Sigma S0819, 1:200).; goat anti-rabbit HRP (Bio-Rad 1705046, 1:10,000); goat anti-mouse HRP (Bio-Rad 1705047, 1:10,000). Target-protein abundance was normalized to vinculin.

### Histological analysis

Unfixed ventricular tissues embedded in optimal cutting temperature compound were stored at -80 °C. Cryosections were cut at a thickness of 6 µm at -16 °C using Leica CM195 cryostat. Hematoxylin and Eosin (H&E) and Picrosirius red (PSR) staining were performed using commercial kits according to the manufacturer’s instructions (Abcam, ab245880 and ab150681, respectively). Whole-section images were acquired by the Rutgers Cellular Imaging and Histology Core using Zeiss Axioscan 7 slide scanner with 20x magnification. Images were analyzed using ImageJ software and the Colour Deconvolution 2 plugin [37, 38]. Mononuclear cell infiltration in H&E-stained sections was quantified using a grid-based scoring method as previously described [14]. Fibrosis in PSR-stained sections was quantified as the percentage of tissue area positive for collagen using a previously described ImageJ-based algorithm [39]. Measurements from technical replicate sections were averaged to generate a single biological value for each mouse.

### Immunofluorescence staining and analysis

Immunofluorescence staining was adapted from our previous protocol [14]. Ventricular cryosections were fixed with 4% paraformaldehyde for 10 minutes at room temperature and washed 3 times in PBS for 5 minutes per wash. Free aldehydes were quenched with 50 mM glycine in PBS for 10 minutes at room temperature, followed by three PBS washes. Sections were permeabilized with 0.1% Triton X-100 in PBS for 5 minutes and washed again. For antibody treatment, sections were blocked for 1 hour at room temperature in PBS containing 1% bovine serum albumin, 5% goat serum, and 0.05% Tween 20. Primary antibodies were applied overnight at 4 °C, followed by fluorophore-conjugated secondary antibodies for 1 hour at room temperature. Wheat germ agglutinin was applied for 10 minutes at room temperature in the dark. Sections were mounted with ProLong Gold antifade reagent with DAPI (Invitrogen, P36935).

Confocal z-stacks consisting of 15 optical sections were acquired using a Leica STELLARIS 8 tau-STED microscope equipped with an HC PL APO CS2 40x/1.25 glycerol-immersion objective. Images were collected using LAS X software with identical acquisition settings for samples included in the same experiment. Image analysis was adapted from our previous study [14]. Z-stack images were processed using the average-intensity method, subjected to rolling-ball background subtraction with a radius of 30 pixels, and converted to 8-bit images in ImageJ. Thresholds were determined using a predefined Otsu algorithm and applied consistently within each experiment. Cx43 localization at the intercalated discs was quantified as the area of Cx43 signal colocalizing with N-cadherin divided by the total Cx43-positive area. Three independently and randomly selected fields were analyzed per mouse, and the resulting values were averaged to generate one biological replicate. The following primary antibodies were used: Cx43 from rabbit (Sigma C6219, 1:200); N-Cadherin (Invitrogen 333900, 1:200); Cx43 from mouse (Invitrogen 138300, 1:200); Nav1.5 (Alomone ASC005, 1:100); goat anti-mouse Alexa 555/647 (Invitrogen A21424/A21235, 1:300); goat anti-rabbit Alexa 555/647 (ThermoFisher A21429/A32733, 1:300); wheat agglutinin-488 (Invitrogen W11261, 1:500).

### Ventricular cardiomyocyte isolation

Adult ventricular cardiomyocytes were isolated by Langendorff perfusion as previously described [14]. Mice received heparin intraperitoneally at 150U per mouse 20 minutes before euthanasia. After deep anesthesia and euthanasia, the heart was excised, cannulated through the aorta, and perfused with Langendorff perfusion buffer (120 mM NaCl, 5.4 mM KCl, 0.3 mM NaH_2_PO_4_, 1.2 mM MgCl_2_, 20 mM HEPES, 10 mM 2,3-butanedione monoxime, 10 mM glucose, 10 mM taurine, pH 7.4). After blood was cleared from the coronary circulation, the heart was perfused for 12 minutes at 37 °C with digestion buffer (310 U/mL collagenase II, Worthington LS004177; 0.104 U/mL protease XIV, Sigma P5147; 30 µM CaCl_2_ in perfusion buffer). The ventricles were subsequently dissected and gently dissociated.

Enzymatic digestion was terminated by adding an equal volume of stop buffer (0.5% BSA, 50 µM CaCl_2_ in perfusion buffer). Cardiomyocytes were enriched through 4 consecutive 10-minute gravity-settling steps. Extracellular calcium was gradually restored using stop buffers containing 0 µM, 100 µM, 400 µM, and 900 µM CaCl_2_. Viable, rod-shaped cardiomyocytes maintained in the final buffer containing 900 µM CaCl_2_ were used for electrophysiological recordings.

### Whole-cell patch clamp recording

The viable cardiomyocytes were initially maintained at 37°C in Normal Tyrode buffer (136 mM NaCl, 5.4 mM KCl, 0.33 mM NaH_2_PO_4_, 1 mM MgCl_2_, 10 mM HEPES, 10 mM glucose, 1 mM CaCl_2_, pH 7.4 by NaOH). Patch pipettes with resistances of 1-3 MΩ were filled with internal solution (70 mM aspartic acid, 60 mM CsCl, 10 mM EGTA, 1 mM MgCl_2_, 10 mM HEPES, 5 mM Na_2_ATP, 1 mM CaCl_2_, pH 7.2 by CsOH) [40]. For sodium-current recordings, the extracellular solution was replaced with Low-Sodium buffer (10 mM NaCl, 5 mM CsCl, 120 mM NMDG, 1.2 mM MgCl_2_, 10 mM HEPES, 5 mM glucose, 10 μM CoCl_2_, 10 μM nifedipine, 1.8 mM CaCl_2_, pH 7.4 by HCl). Whole-cell currents were recorded in voltage-clamp mode using MultiClamp 700A amplifier (Axon Instruments) and pCLAMP software (Version 10.2.0.12, Axon Instruments). Cells were held at -90 mV and depolarized from -80 to +50 mV with 5 mV increments. Series resistance was compensated by 40% to 60% using the amplifier circuitry to minimize voltage-clamp error. Currents were normalized to whole-cell capacitance and expressed as current density. Recordings with excessive leakage, unstable access resistance, prominent voltage-clamp artifacts, or nonexponential capacitive-current decay were excluded from analysis. Data were analyzed using Clampfit software (version 11.2.2.17).

### Statistics

Statistical analyses were performed using GraphPad Prism. Data are presented as mean ± SEM. The mouse was considered the experimental unit for in vivo, molecular, biochemical, and histological experiments. For patch clamp experiments, both the number of cells analyzed (n) and the number of mice from which the cells were obtained (N) are reported in the figure legends. For experiments involving genotype and treatment as independent factors, data were analyzed using ordinary 2-way ANOVA followed by the Holm-Šídák’s multiple-comparisons procedure. Prespecified comparisons were made between cell means within the same genotype or treatment category, as indicated in the corresponding figure legends. Other statistical tests, when applicable, are specified in the figure legends. A 2-sided P value < 0.05 was considered statistically significant. All quantification and analysis were performed with investigators blinded to genotype and treatment.

## Study approval

The animal studies were approved by the Institutional Animal Care and Use Committee (IACUC) of Rutgers New Jersey Medical School and conducted in accordance with NIH guidelines. All the tissue samples were collected postmortem.

## Data and Materials Availability

The data that support the findings of this study are available from the corresponding author upon reasonable request. The availability of genetically modified mouse lines and detailed viral vector design information is subject to institutional approval and applicable material transfer agreements.

## AI disclosure

Generative AI-assistance was used only for grammar, spelling, and language editing during manuscript preparation.

## Authorship contribution

**D. Z.**: Writing (original draft, review, editing), Visualization, Validation, Software, Methodology, Investigation, Funding acquisition, Formal analysis, Data curation, Conceptualization.

**V. Y.**: Validation, Software, Investigation, Formal analysis, Data curation.

**N. K. A.**: Investigation, Software, Data curation.

**G. G.**: Investigation, Data curation.

**E. M.**: Investigation, Software, Data curation.

**L. H. X.**: Writing (review, editing), Resources, Methodology, Funding acquisition, Formal analysis.

**D. F.**: Writing (review, editing), Visualization, Validation, Supervision, Resources, Project administration, Methodology, Investigation, Funding acquisition, Formal analysis, Data curation, Conceptualization.

## Funding

This work was supported by an American Heart Association (AHA) Predoctoral Fellowship 23PRE1019365 (to D.Z.); NIH grant R01HL171094 (to D.F.); and NIH grant R01HL157116 (to L.H.X.). The confocal images were obtained from a Leica TCS SP8 STED 3x super resolution microscope, funded by NIH grants 1S10OD025182-01A1 for the support of the Rutgers Cellular Imaging and Histology Core (to project leader Dongfang Liu, RBHS Newark campus).

## Declaration of competing interest

None

## Acknowledgements

We thank Dr. Luke Fritzky, Joel Pierre, and Joey Bulatowicz (Rutgers University) for assistance in histology and microscopy. We thank Dr. Glenn I. Fishman (New York University) for providing original Cx43-S3E founder mice for our colonies. We thank Dr. Ghassan Yehia and Dr. Peter J. Romanienko (Rutgers Robert Wood Johnson Medical School) for generating the original Tubb3-S172E founder mice for our colonies. We thank VectorBuilder for assistance and packaging the AAV9 vectors. The graphical abstract was created with BioRender.com.

---

**Supplemental Figure 1.**
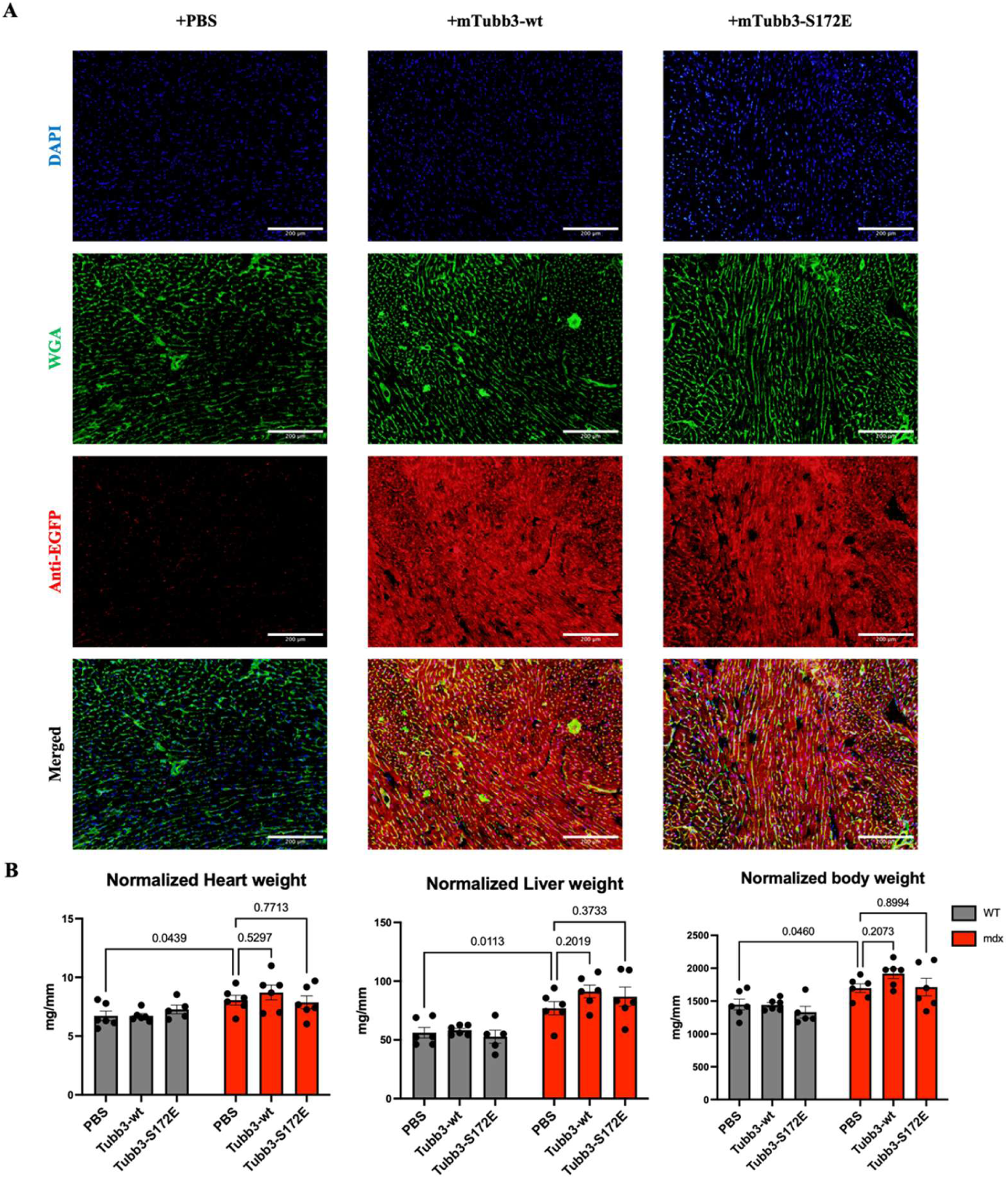
Validation and gross physiological assessment of AAV9-treated mice. **(A)** Representative immunofluorescence images of ventricular sections from mdx mice treated with PBS, AAV9-mTubb3-WT, or AAV9-mTubb3-S172E. Direct EGFP fluorescence was not readily detectable in the 488 nm channel; therefore, transgene reporter expression was assessed by anti-EGFP immunostaining. EGFP immunofluorescence was detected in AAV9-treated hearts and showed the expected expression. Sections were counterstained with WGA-Alexa 488 to visualize tissue architecture. Immunofluorescence staining was performed on OCT-embedded cryosections as described in the Methods with following antibodies: anti-EGFP (Abcam ab6556, 1:200), WGA-Alexa 488 (Invitrogen W11261, 1:500), goat anti-rabbit Alexa 555 (ThermoFisher A21429, 1:300). Images were acquired using Olympus BX51 microscope with UPlanApo 10x/0.40 objective. Scale bar = 200 µm. **(B)** Heart weight, liver weight, and body weight normalized to tibia length across experimental groups. Data are presented as mean ± SEM. Sample size was n=6 for each group, except for WT mice treated with AAV9-Tubb3-S172E, where n=5. Statistical analysis was performed using two-way ANOVA followed by Holm-Sidak post hoc multiple-comparisons test. Exact P values are indicated in the figure.

**Supplemental Figure 2.**
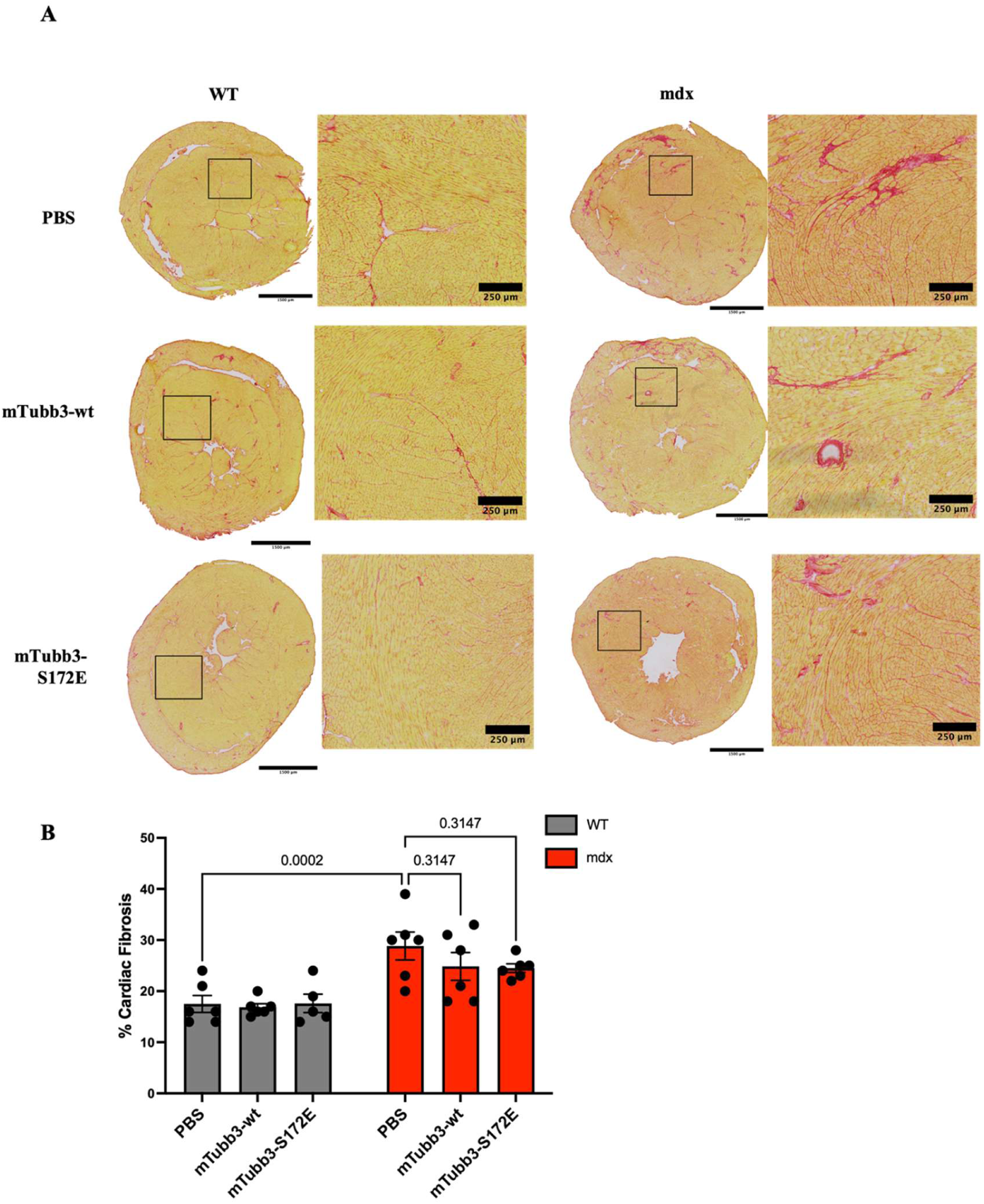
Assessment of cardiac fibrosis following AAV9-mediated expression of WT or phospho-mimic S172E βIII-tubulin. **(A)** Representative Picrosirius Red (PSR)-stained ventricular sections from WT and mdx mice treated with PBS, AAV9-mTubb3-WT, or AAV9-mTubb3-S172E at 4-5 months of age. Higher-magnification insets show representative regions of collagen deposition. **(B)** Quantification of cardiac fibrosis, expressed as the percentage of PSR-positive area relative to the total tissue area. Scale bars: 1500 μm for low-magnification images and 250 μm for higher-magnification insets. Data are presented as mean ± SEM. Sample size was n=6 for each group, except for WT mice treated with AAV9-Tubb3-S172E, where n=5. Statistical analysis was performed using two-way ANOVA followed by Holm-Sidak post hoc multiple-comparisons test. Exact P values are indicated in the figure.

## Uncut Western blot

### Figure 1C and 5A

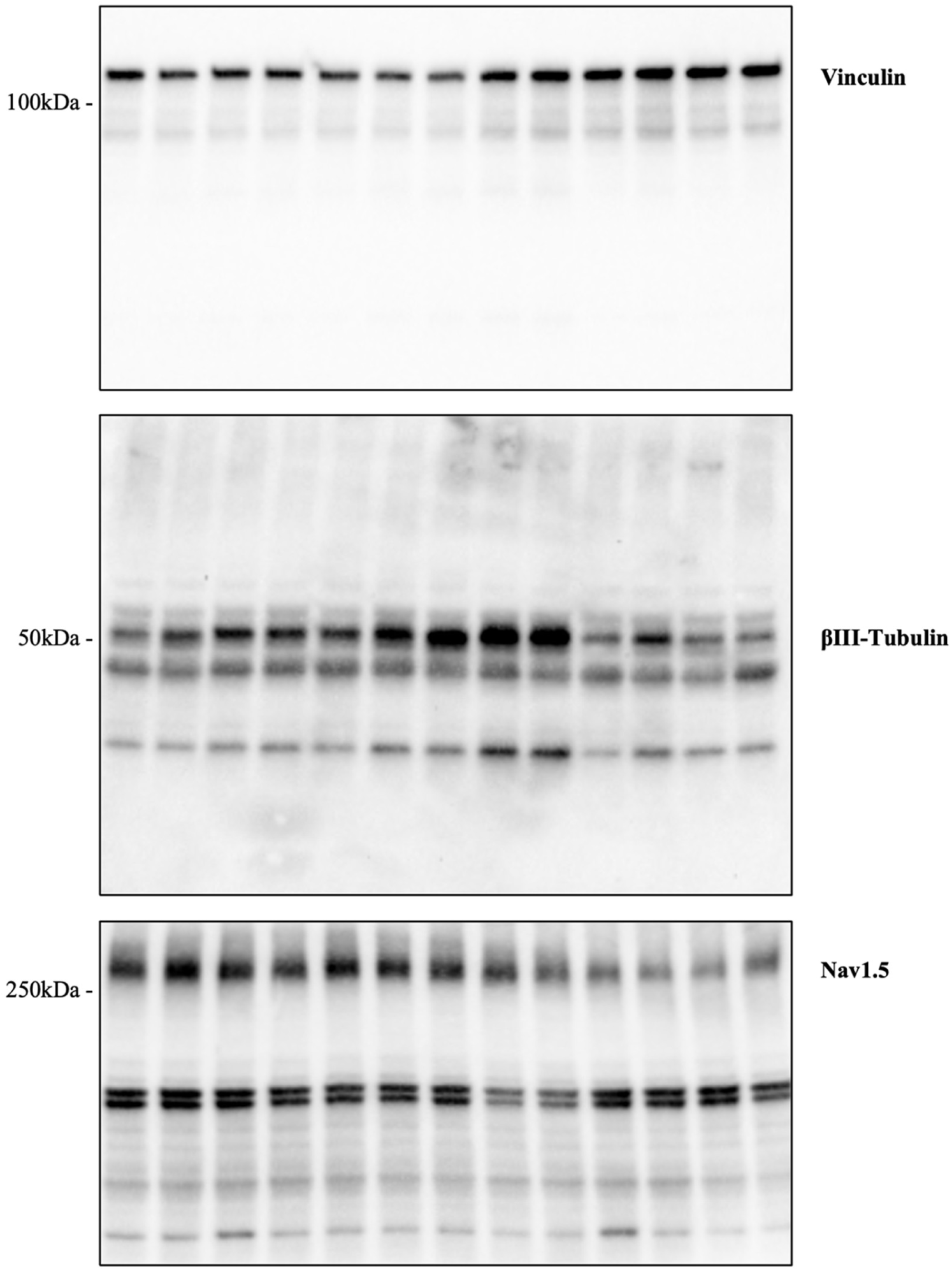

## Notes

### Competing Interest Statement

The authors have declared no competing interest.

## References

1. Lechner, A., et al., Cardiomyopathy as cause of death in Duchenne muscular dystrophy: a longitudinal observational study. ERJ Open Res, 2023. 9(5): p. 00176–2023.

2. Schultz, T.I., F.J. Raucci, and F.N. Salloum, Cardiovascular Disease in Duchenne Muscular Dystrophy. JACC: Basic to Translational Science, 2022. 7(6): p. 608–625.

3. Tandon, A., et al., Myocardial Fibrosis Burden Predicts Left Ventricular Ejection Fraction and Is Associated With Age and Steroid Treatment Duration in Duchenne Muscular Dystrophy. Journal of the American Heart Association, 2015. 4(4): p. e001338.

4. Wilson, D.G.S., A. Tinker, and T. Iskratsch, The role of the dystrophin glycoprotein complex in muscle cell mechanotransduction. Commun Biol, 2022. 5(1): p. 1022.

5. Lapidos, K.A., R. Kakkar, and E.M. McNally, The Dystrophin Glycoprotein Complex. Circulation Research, 2004. 94(8): p. 1023–1031.

6. Uchida, K., E.A. Scarborough, and B.L. Prosser, Cardiomyocyte Microtubules: Control of Mechanics, Transport, and Remodeling. Annu Rev Physiol, 2022. 84: p. 257–283.

7. Caporizzo, M.A., C.Y. Chen, and B.L. Prosser, Cardiac microtubules in health and heart disease. Exp Biol Med (Maywood), 2019. 244(15): p. 1255–1272.

8. Warner, E.F., Y. Li, and X. Li, Targeting Microtubules for the Treatment of Heart Disease. Circulation Research, 2022. 130(11): p. 1723–1741.

9. Nasilli, G., et al., Decreasing microtubule detyrosination modulates Nav1.5 subcellular distribution and restores sodium current in mdx cardiomyocytes. Cardiovasc Res, 2024. 120(7): p. 723–734.

10. Kerr, J.P., et al., Detyrosinated microtubules modulate mechanotransduction in heart and skeletal muscle. Nature Communications, 2015. 6(1): p. 8526.

11. Prins, K.W., et al., Microtubule-Mediated Misregulation of Junctophilin-2 Underlies T-Tubule Disruptions and Calcium Mishandling in mdx Mice. JACC Basic Transl Sci, 2016. 1(3): p. 122–130.

12. Himelman, E., et al., A microtubule-connexin-43 regulatory link suppresses arrhythmias and cardiac fibrosis in Duchenne muscular dystrophy mice. American Journal of Physiology-Heart and Circulatory Physiology, 2022. 323(5): p. H983–H995.

13. Himelman, E., et al., Prevention of connexin-43 remodeling protects against Duchenne muscular dystrophy cardiomyopathy. Journal of Clinical Investigation, 2020. 130(4): p. 1713–1727.

14. Zhou, D., et al., Phospho-mimic βIII-tubulin rescues microtubule and cardiac defects in Duchenne muscular dystrophy mice. Journal of Molecular and Cellular Cardiology, 2026. 215: p. 6–19.

15. Boengler, K., et al., Importance of Cx43 for Right Ventricular Function. Int J Mol Sci, 2021. 22(3): p. 987.

16. Severs, N.J., et al., Remodelling of gap junctions and connexin expression in diseased myocardium. Cardiovascular Research, 2008. 80(1): p. 9–19.

17. Lillo, M.A., et al., S-nitrosylation of connexin43 hemichannels elicits cardiac stress–induced arrhythmias in Duchenne muscular dystrophy mice. JCI insight, 2019. 4(24): p. e130091.

18. Araujo, P.A., et al., Connexin-43 remodelling and arrhythmias: hemichannels as key drivers of cardiac dysfunction. J Physiol, 2025. 603(15): p. 4293–4306.

19. Lillo, M.A., et al., Remodeled connexin 43 hemichannels alter cardiac excitability and promote arrhythmias. Journal of General Physiology, 2023. 155(7).

20. Gonzalez, J.P., et al., Normalization of connexin 43 protein levels prevents cellular and functional signs of dystrophic cardiomyopathy in mice. Neuromuscul Disord, 2018. 28(4): p. 361–372.

21. Le Dour, C., et al., Actin-microtubule cytoskeletal interplay mediated by MRTF-A/SRF signaling promotes dilated cardiomyopathy caused by LMNA mutations. Nature Communications, 2022. 13(1).

22. Yu, X., et al., MARK4 controls ischaemic heart failure through microtubule detyrosination. Nature, 2021. 594(7864): p. 560–565.

23. Chen, C.Y., et al., Suppression of detyrosinated microtubules improves cardiomyocyte function in human heart failure. Nature Medicine, 2018. 24(8): p. 1225–1233.

24. Caporizzo, M.A., et al., Microtubules Increase Diastolic Stiffness in Failing Human Cardiomyocytes and Myocardium. Circulation, 2020. 141(11): p. 902–915.

25. Khairallah, R.J., et al., Microtubules underlie dysfunction in duchenne muscular dystrophy. Sci Signal, 2012. 5(236): p. ra56.

26. Janke, C. and M.M. Magiera, The tubulin code and its role in controlling microtubule properties and functions. Nature Reviews Molecular Cell Biology, 2020. 21(6): p. 307–326.

27. Pachter, J.S., T.J. Yen, and D.W. Cleveland, Autoregulation of tubulin expression is achieved through specific degradation of polysomal tubulin mRNAs. Cell, 1987. 51(2): p. 283–292.

28. Lin, Z., et al., TTC5 mediates autoregulation of tubulin via mRNA degradation. Science, 2020. 367(6473): p. 100–104.

29. Ohi, R., C. Strothman, and M. Zanic, Impact of the ’tubulin economy’ on the formation and function of the microtubule cytoskeleton. Curr Opin Cell Biol, 2021. 68: p. 81–89.

30. Frangogiannis, N.G., Cardiac fibrosis: Cell biological mechanisms, molecular pathways and therapeutic opportunities. Mol Aspects Med, 2019. 65: p. 70–99.

31. Phyo, S.A., et al., Transcriptional, post-transcriptional, and post-translational mechanisms rewrite the tubulin code during cardiac hypertrophy and failure. Frontiers in Cell and Developmental Biology, 2022. 10: p. 837486.

32. Shaw, R.M., et al., Microtubule plus-end-tracking proteins target gap junctions directly from the cell interior to adherens junctions. Cell, 2007. 128(3): p. 547–60.

33. Meyers, T.A. and D. Townsend, Cardiac Pathophysiology and the Future of Cardiac Therapies in Duchenne Muscular Dystrophy. Int J Mol Sci, 2019. 20(17): p. 4098.

34. Leo-Macias, A., E. Agullo-Pascual, and M. Delmar, The cardiac connexome: Non-canonical functions of connexin43 and their role in cardiac arrhythmias. Semin Cell Dev Biol, 2016. 50: p. 13–21.

35. Stroemlund, L.W., et al., Gap junctions - guards of excitability. Biochem Soc Trans, 2015. 43(3): p. 508–12.

36. Livak, K.J. and T.D. Schmittgen, Analysis of relative gene expression data using real-time quantitative PCR and the 2(-Delta Delta C(T)) Method. Methods, 2001. 25(4): p. 402–8.

37. Ruifrok, A.C. and D.A. Johnston, Quantification of histochemical staining by color deconvolution. Anal Quant Cytol Histol, 2001. 23(4): p. 291–9.

38. Landini, G., G. Martinelli, and F. Piccinini, Colour deconvolution: stain unmixing in histological imaging. Bioinformatics, 2021. 37(10): p. 1485–1487.

39. Hildyard, J.C.W., et al., Rapid histological quantification of muscle fibrosis and lysosomal activity using the HSB colour space. bioRxiv, 2022: p. 2022.08.02.502489.

40. Albesa, M., et al., Regulation of the cardiac sodium channel Nav1. 5 by utrophin in dystrophin-deficient mice. Cardiovascular research, 2011. 89(2): p. 320–328.

